# Visual cortical entrainment to unheard acoustic speech reflects intelligibility of lip movements and is mediated by dorsal stream regions

**DOI:** 10.1101/244277

**Authors:** A. Hauswald, C. Lithari, O. Collignon, E. Leonardelli, N. Weisz

**Author notes:** These authors contributed equally (A. Hauswald, C. Lithari).

## Abstract

Successful lip reading requires a mapping from visual to phonological information [1]. Recently, visual and motor cortices have been implicated in tracking lip movements (e.g. [2]). It remains unclear, however, whether visuo-phonological mapping occurs already at the level of the visual cortex, that is, whether this structure tracks the acoustic signal in a functionally relevant manner. In order to elucidate this, we investigated how the cortex tracks (i.e. entrains) *absent* acoustic speech signals carried by silent lip movements. Crucially, we contrasted the entrainment to unheard forward (intelligible) and backward (unintelligible) acoustic speech. We observed that the visual cortex exhibited stronger entrainment to the unheard forward acoustic speech envelope compared to the unheard backward acoustic speech envelope. Supporting the notion of a visuo-phonological mapping process, this forward-backward difference of occipital entrainment was not present for actually observed lip movements. Importantly, the respective occipital region received more top-down input especially from left premotor, primary motor, somatosensory regions and, to a lesser extent, also from posterior temporal cortex. Strikingly, across participants, the extent of top-down modulation of visual cortex stemming from these regions partially correlates with the strength of entrainment to absent acoustic forward speech envelope but not to present forward lip movements. Our findings demonstrate that a distributed cortical network, including key dorsal stream auditory regions [3–5], influence how the visual cortex shows sensitivity to the intelligibility of speech while tracking silent lip movements.

**Highlights:**

- Visual cortex tracks better forward than backward unheard acoustic speech envelope
- Effects not “trivially” caused by correlation of visual with acoustic signal
- Stronger top-down control of visual cortex during forward display of lip movements
- Top-down influence correlates with visual cortical entrainment effect
- Results seem to reflect visuo-phonological mapping processes

## Results

Successful lip reading in absence of acoustic information requires mechanisms of mapping from the visual information to the corresponding but absent phonological code [1]. We know that the visual and motor cortices track lip movements for congruent compared to incongruent audio-visual speech [2], but the large-scale neural processes precisely involved in linking visual speech (lip movements) processing with the auditory content of the speech signal has remained obscure. We performed a Magnetoencephalography (MEG) experiment in which 24 participants were exposed to silent lip movements corresponding to forward and backward speech. In a parallel behavioral experiment with 19 participants, we demonstrate that silent forward lip movements are intelligible while backward presentation is not: participants could correctly identify words above chance level when presented with silent forward visual speech, while performance for silent backward visual speech did not differ from chance level. We neurally compared the tracking of the unheard acoustic speech by contrasting the coherence between (unheard) forward and backward acoustic speech envelope with the brain activity elicited by silently presented forward and backward lip movements. Uncovering visual cortical regions via this analysis, we then performed Granger causality analysis to identify the cortical regions mediating top-down control and assessed to what extent this was correlated to the aforementioned entrainment effect. Importantly, we also analyzed occipital forward-backward entrainment and Granger causality for coherence between lip contour and brain activity during visual speech to show that the findings are specific for the envelope of (unheard) acoustic speech-brain coherence.

### Audiovisual coherence peaks at 5 Hz for forward and backward presentations

A short example of the lip signal and the corresponding audio signal as well as its envelope is depicted in Fig. 1A. The coherence spectrum between lip contour and acoustic speech – lip-speech coherence – (Fig. 1B) exhibits a distinct peak at 5 Hz, matching well the syllable rhythm of speech [6,7]. Contrasting forward and backward lip-speech coherence for the whole frequency band (1–20 Hz) did not reveal any differences (all *p*’s > .05). This excludes attributing differences between forward and backward coherences of acoustic speech envelope with the brain signal to low level differences in the nature of forward and backward acoustic speech envelope and lip contour.

**Fig. 1.**
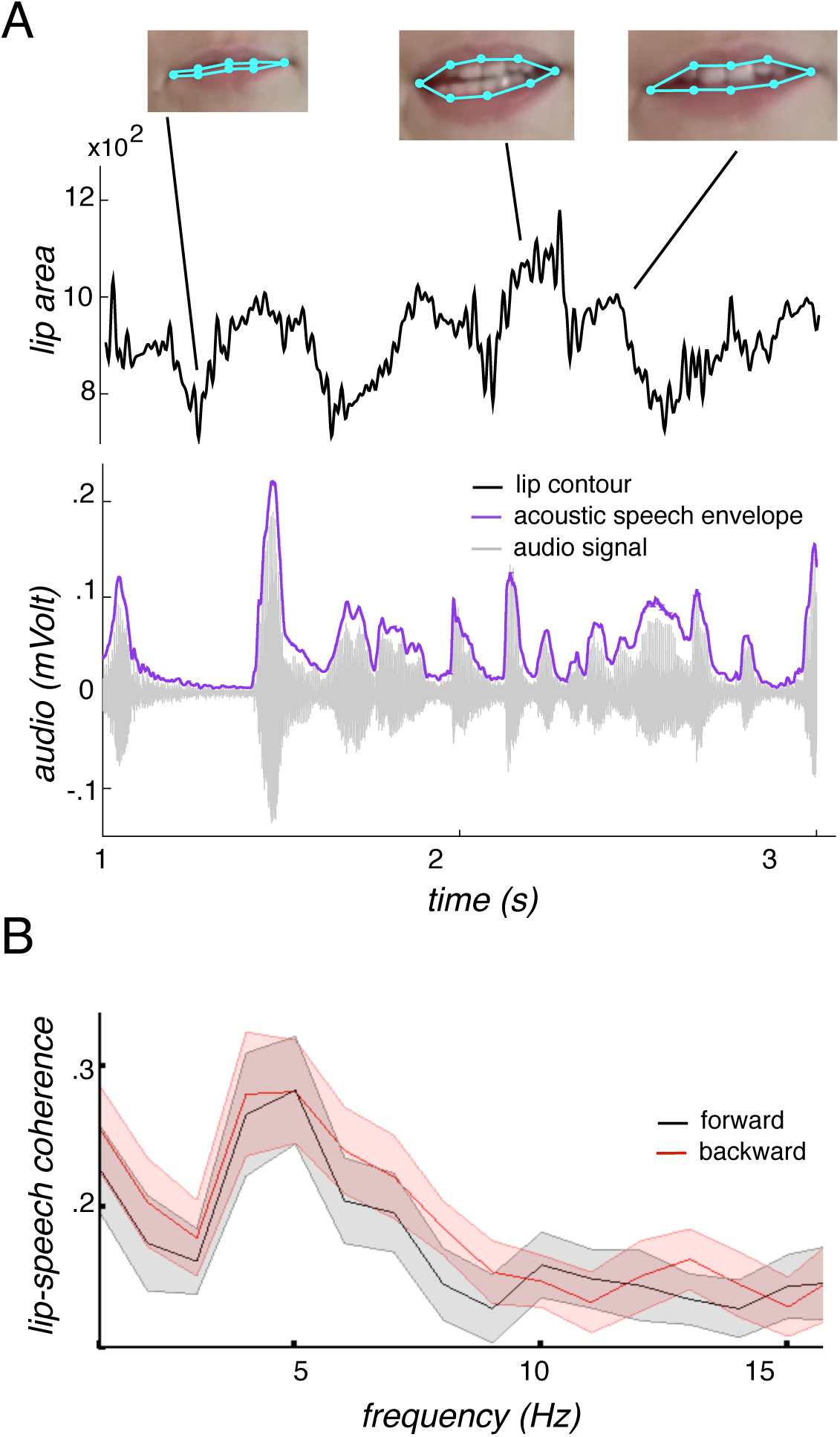
Lip contour and acoustic speech envelope. **A)** The time series of the lip contour (area defined by the light blue points measured in square pixels) with the corresponding video frames is displayed together with the audio signal and the acoustic speech envelope [8] for a 3 second forward section. For the coherence analyses, the lip contour and the acoustic speech envelope were used. **B)** The coherence spectrum (averaged across stimuli) between lip contour and acoustic speech envelope is plotted peaking at 5 Hz, the rhythm of the syllables, for both forward and backward speech. The shaded area reflects the standard deviation across stimuli. No difference between forward and backward speech occurred.

### Forward presentation of silent lip movements more intelligible than backward presentations

To ensure that silent lip movements differ in terms of intelligibility when presented forwards or backwards, we performed a separate behavioral experiment. Participants watched short videos of silent visual speech and were then asked to choose between two words, one of which was contained in the short video. Hits (mean: 64.84%) in the forward condition were significantly higher than chance level (*t*(18)=7.81, *p*<0.0005) while this was not the case for hits in the backward condition (mean: 53.47%, *t*(18)=1.54, *p*=0.14). Hits in the forward condition were also significantly higher than hits in the backward condition (*t*(18)=3.76, *p*<.005; Fig. 2A).

**Fig. 2:**
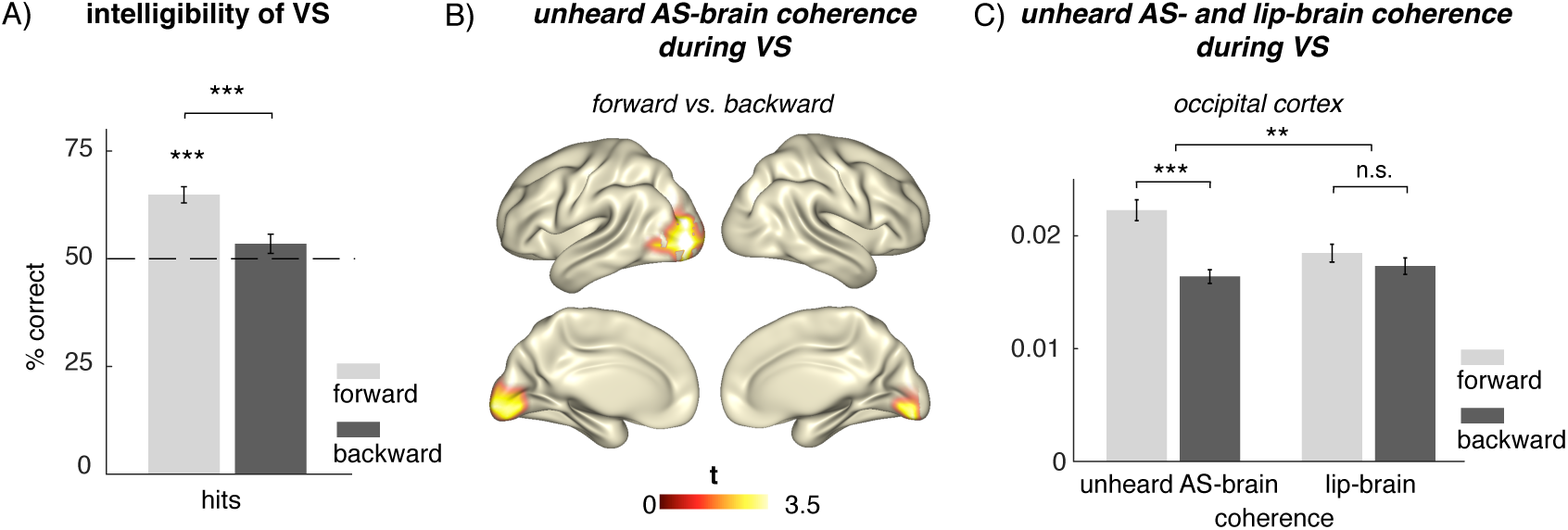
**A)** Separate behavioral experiment on intelligibility in silent visual speech (VS) with 19 participants. The contrast hits vs. chance level was significant only in the forward condition. Hits in the forward condition were also higher than hits in the backward condition. **B)** Coherences of theta band brain sources (4–7 Hz) with the not-heard acoustic envelope of speech (AS, while watching visual speech, VS, contrasted between forward and backward conditions, *p* < .05, Monte-Carlo corrected) is increased at occipital regions when watching lip movements of forward speech compared to lip movements of backward speech. **C)** Mean of the individual unheard acoustic speech-brain and lip-brain coherence values during forward and backward presentation of visual speech extracted at the voxels of the statistical effect found in B. Difference in occipital cortex between forward and backward unheard acoustic speech-brain coherence bigger than the lip-brain coherence during visual speech (n.s.). Error bars indicate standard error. ** p<.005, *** p < .0005.

### Stronger entrainment between occipital activity and the envelope of acoustic speech during forward presentation of lip movements

We first calculated the coherence between the absent acoustic envelope of the speech signal with the source reconstructed *(LCMV)* MEG data on each voxel while participants were watching the silent lip movements. Occipital regions showed a statistically significant difference between the neural response to forward versus backward unheard acoustic speech envelope (*t*(23)=6.83, *p*<.000005, Fig. 2B and C). Given the high coherence between the acoustic and the visual signal related to speech stimuli (see above), the aforementioned effect could be a trivial by-product of a differential entrainment to the visual information only. To test this possibility, we investigated the coherence between occipital activity and the lip contour (lip-brain coherence). The forward-backward difference was bigger for the unheard acoustic speech-brain coherence carried by lip movements (*t*(23)=3.41, *p*<0.005) than for lip-brain coherence elicited by lip movements (*t*(23)=0.97, *p*=0.34; Fig. 2C). This implies that the visual cortex tracks lip movements faithfully regardless of whether they are displayed forwards or backwards. However, only forward presented lip movements additionally elicit an entrainment of visual cortical activity to the envelope of the corresponding (unplayed) acoustic signal.

### Top-down modulation on visual cortex drives (unheard) acoustic speech envelope entrainment effects

In order to elucidate the network driving this putative visuo-phonological mapping process, we calculated Granger causality between the occipital region showing the strongest difference between forward and backward acoustic speech entrainment and the remaining whole-brain voxels. We focused the statistical analyses on relevant regions (parietal, temporal, pre-and postcentral areas in the left hemisphere) that include regions of interest as motivated by dual-route models of speech processing [3–5]. The contrast (corrected for multiple comparisons; see methods) between Granger causality for forward and backward visual speech revealed a positive effect in left premotor, primary motor and primary somatosensory cortex (Fig. 3A). At a descriptive level, also posterior portions of the left superior temporal cortex (BA 22), inferior temporal gyrus (BA 37), and middle temporal gyrus (BA 39 including the angular gyrus) were above the uncorrected statistical critical value (see methods). Altogether, this means that key nodes of mainly dorsal route processing regions exert relatively more top-down influence on the visual cortex during forward compared to backward presentation of visual speech.

**Fig. 3:**
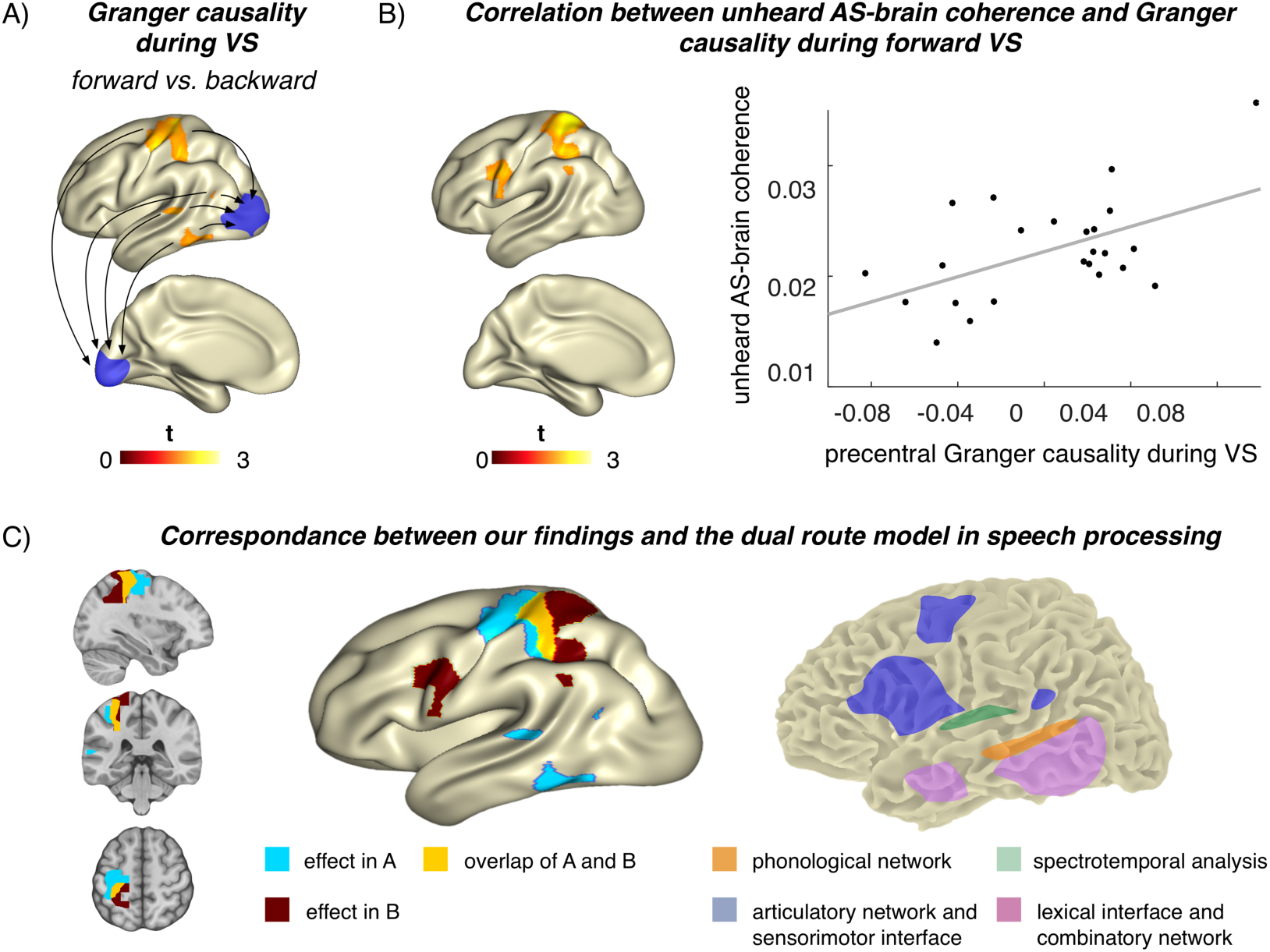
**A)** Granger causality during forward vs. backward lip movements (visual speech, VS) using occipital seed region taken from an effect found for the envelope of unheard acoustic speech (AS)-brain coherence (blue) that was defined as voxel with the strongest forward-backward entrainment effect to unheard acoustic speech envelope. Calculation of contrast between normalized ((ingoing − outgoing)/(ingoing + outgoing)) forward and backward visual speech shows positive effects in left premotor, primary motor and primary sensory cortex (corrected for multiple comparisons) and (uncorrected) posterior superior (BA 22) and inferior temporal gyrus (BA 37) as well as the posterior middle temporal gyrus including the angular gyrus (BA 39). Maps are masked by uncorrected statistical critical value (1.714). **B)** Significant correlation between the occipital unheard acoustic speech-brain coherence and the Granger causality of forward visual speech. **Left**: Corrected for multiple comparison in pre-/postcentral gyrus. Also, above the uncorrected statistical critical value were inferior premotor cortex (BA 6), frontal eye fields (BA 8), and posterior middle temporal gyrus (BA 39). **Right**: Scatterplot of correlation (*r*=0.46, *p*<0.05) in pre-/postcentral gyrus. **C)** Illustration of the correspondence between our findings and the proposed dual route model of speech processing [5]. **Left:** Illustration of the effects above the uncorrected statistical critical value found for differences in Granger causality between forward and backward visual speech (blue, see also Fig. 3A), correlation between unheard acoustic speech-brain coherence and Granger causality of forward visual speech (Fig., 3B, red) and their overlap (yellow) in anatomical and surface view. **Right**: Illustration of the dual route model as proposed by Hickock and Poeppel (adapted from [5]) comprising the dorsal route (blue) and the ventral route (red).

To clarify whether the network level effect was actually related to the unheard acoustic speech-brain coherence or a mere by-product of differential lip-brain coherence, we calculated the correlations of the forward Granger causality with both forward acoustic speech-brain coherence and lip-brain coherence in occipital regions. Only the correlations with unheard acoustic speech-brain coherence revealed significant results while the lip-brain coherence did not (*p*>0.4). For the unheard acoustic speech-brain coherence, mainly precentral and postcentral regions revealed a strong correlation (corrected for multiple comparisons, *p*<.05). At an uncorrected level the premotor (BA 6), frontal eye field (BA 8), and posterior middle temporal regions (BA 39) also yielded correlations above the statistical critical value (see. Fig. 3B, left). The scatterplot in Fig. 3B (right) illustrates this correlation for a precentral region (*r*=0.46, *p*<0.05) that showed a statistical effect for the forward-backward Granger causality contrast during visual speech (Fig. 3A) and the correlation between unheard acoustic speech-brain coherence and Granger causality (Fig. 3B, left). Interestingly these findings show high correspondence with the relevant regions of the dual-route model of speech [5] (see Fig. 3C).

## Discussion

Dual-process models of speech processing state the presence of two different routes [3–5]. While the ventral stream is assumed to contribute to comprehension, the dorsal stream is proposed to “map acoustic speech signals to frontal lobe articulation” [5]. Extending this notion Rauschecker [3] proposes the dorsal stream to be a “supramodal reference frame”, enabling flexible transformation of sensory signals (see also [9]). Most evidence underlying the development of these models used acoustic signals (e.g. [10,11]).

Already very early in life, an audio-visual link between observing lip movements and hearing speech sounds is present, which consequentially supports infants to acquire their first language. In this context, visual speech constitutes a crucial role for speech processing, as can be seen by findings of infants of just 4 to 6 month of age who can discriminate their native language from silent lip movements only [12] or who allocate more attention to the mouth compared to the eyes once they detect synchrony between lip movements and speech sounds [13]. The importance of lip reading for speech processing is also demonstrated by studies with deaf individuals [14], showing that lip-reading alone can be sufficient for language comprehension suggesting a functional role that goes beyond the mere support of auditory input processing. In a recent study, Lazard and Giraud [1] hypothesized that the functional role of lip reading was to enable the visual cortex to remap visual information into phonological information. Given the described mapping processes, we ask the following question: how does the mapping of an acoustic signal influence lip reading when the acoustic signal is absent and only induced by the lip movements. We investigated this via cortical entrainment to the absent acoustic speech signal that was carried by the silent lip movements. We contrasted forward and backward visual conditions of speech segments, given that only the forward condition was intelligible as shown by the behavioral experiment and therefore should induce speech-related processes.

The auditory cortex has been repeatedly found to be activated and entrained in response to audio-only [15] or audio-visual stimulation [16,17]. Previous fMRI studies established also that perception of silent visual speech activates the auditory cortices, clearly showing involvement of auditory processes during lip-reading [18,19]. Recently, a study attempted a reconstruction of posterior surface EEG channels (presumably capturing visual cortex activity) via the absent acoustic speech envelope [20]. The authors showed that a model based on the absent acoustic signal predicted posterior EEG activity similar to models based on frame-to-frame motion changes and categorical visual speech features [20]. However, since the acoustic signal was seen as a proxy for lip movements (which were not explicitly investigated), the separate contributions of acoustic and visual information were not explored. Going beyond this finding, we investigated cortical entrainment to the absent acoustic signal by comparing forward and backward envelopes of unheard acoustic speech carried by silent presentations of lip movements putatively linked to altered speech intelligibility (see Fig. 2A). Using source level analysis, we found that the visual cortex showed stronger entrainment (higher coherence) to unheard acoustic speech envelope during forward (intelligible) rather than backward (unintelligible) mute lip movements. Importantly, our control analysis of coherence between brain activity and the lip contour of the actually observed lip movements did not reveal similar forward vs backward differences (Fig. 2C). This excludes the possibility that our findings for unheard acoustic speech-brain coherence are just an epiphenomenon, given that lip and audio signals are highly correlated (Fig. 1B; cf. [21]). Further, this control analysis argues against the possibility that attentional processes produced the difference in forward-backward unheard acoustic speech-brain coherence, as for both measures (unheard acoustic speech-brain coherence and lip-brain coherence) identical datasets of MEG recordings were used. To our knowledge, this is the first time that an effect of speech intelligibility on brain coherence with unheard acoustic speech has been reported.

It was recently shown that the visual cortex is important for tracking speech related information, for example sign language [22] and lip movements [2]. Further, the more adverse the condition (low signal-to-noise-ratio), the more the visual cortex is entrained to the speech signal of actual acoustic speech presented together with varying levels of acoustic noise and either informative or uninformative visual lip movements at low frequencies [21]. Our study confirms and extends these findings by investigating how occipital regions track the acoustic envelope related to silent visual speech delivered by lip movements. We show that in the absence of acoustic information, the unheard auditory signal – not the lip movement – entrains the visual cortex differentially for intelligible and unintelligible speech. In this case, a visuo-phonological mapping mechanism needs to be in place, and our results showing entrainment of visual area to (non-presented) acoustic speech may be a reflection of such a process. This mapping presumably interacts with top-down processes as lip-reading has been reported to benefit from contextual cues [23]. Recent studies provide evidence for top-down processes in audio-visual and audio-only settings. For example, Giordano and colleagues (2017) showed an increase in directed connectivity between superior frontal regions and visual cortex under the most challenging (acoustic noise and uninformative visual cues) conditions. Kayser et al. [24] proposed top-down processes modulating acoustic entrainment. Park et al. [25] showed enhanced top-down coupling between frontal and, in their case, auditory (due to only auditory stimuli) regions during intelligible speech in the left hemisphere compared to unintelligible speech. Going one step further, given the complete absence of auditory input during silent visual speech, we also expected similar enhanced top-down control to differentiate intelligibility but, in our case, of the visual cortex. Indeed, calculating Granger causality (4–7 Hz, forward and backward visual speech) between visual regions and the other regions of interest showed differential ingoing and outgoing connections for the two conditions: We contrasted the two normalized [(ingoing−outgoing)/(ingoing+outgoing)] conditions and, as expected, the forward condition yielded a more positive ratio of ingoing and outgoing connections than the backward condition stemming from mainly left (pre-)motor regions and primary sensory regions. Also in auditory or audio-visual studies [2,21,24,25], motor regions play an important role in top-down control.

Further, the posterior portions of left superior, middle, and inferior temporal gyrus show differences in the statistical comparison although not significant at a cluster-corrected level. They are nevertheless reported because they provide interesting insights given their previously established role in speech processing [10], particularly under adverse conditions [26] and audiovisual integration, respectively (overview in [27,28]).

Importantly, the significance of the top-down processes for visuo-phonological mapping is further supported by the correlations between the acoustic speech-brain coherence in occipital regions during silent visual speech and the Granger causality effect. We find positive correlations mainly for precentral and postcentral regions but also at premotor areas (BA 6), the frontal eye field (BA 8), and the posterior middle temporal gyrus (BA 39). The precentral and postcentral areas showed a strong overlap with the regions that had enhanced Granger causality in the forward condition. The positive correlations in these regions suggest that the extent of top-down influence on visual cortical regions is associated, at least partially, with the magnitude of this region to exhibit entrainment to (not presented) acoustic speech input. Importantly, the Granger causality effects are not a by-product of the entrainment to the lip movements, as shown by the missing correlations between occipital lip-brain coherence and Granger causality in the regions of interest (relevant for the dual route model).

## Conclusions

Our findings suggest that while observing lip movements, acoustic speech entrains the visual cortex even if the auditory part of the speech input is physically not present. Importantly this cortical mechanism is sensitive to intelligibility, while the same is not the case when looking at entrainment to the actual visual signal. Thus, while observing forward (and more intelligible) lip movements, visual cortex additionally tracks the absent acoustic envelope. Our results strongly suggest dorsal stream regions, including motor-related areas, may mediate this visuo-phonological mapping process by exerting top-down control of visual cortex.

Referring again to the initially mentioned dual-route model of speech processing [3–5], our results show a strikingly high correspondence of involved regions. This underlines the importance of these regions in processing speech relevant information across modalities. Overall, our study supports the idea that in particular dorsal processing routes are activated by observation of silent lip movements, enabling a top-down controlled mapping of the visual signal into the absent acoustic signals [1]. This mapping might be achieved via functional dependencies between auditory and visual sensory systems that have existed since the earliest stages of sensory processing in humans [29–31].

## Acknowledgements

The authors would like to thank Dr. Markus Van Ackeren for his help in extracting the speech envelope. The contribution of CL and NW was financed by the European Research Council (WIN2CON, ERC, StG 283404). OC is supported by the European Research Council (MADVIS, ERC, StG 337573)

## Author Contributions

N.W. and O.C. designed the experiment; E.L. ran the behavioral experiment. A.H. and C.L. analysed the data. All authors wrote the paper.

## Declaration of Interests

The authors declare no competing interests.

## Methods

### Contact for reagent and resource sharing

Further information and requests for resources and reagents should be directed to and will be fulfilled by the Lead Contact, Nathan Weisz.

### Experimental model and subject details

Twenty-three healthy volunteers (age 28.31 ± 4.6, 9 females, all right handed) with normal hearing and vision participated in the study. All participants were native Italian speakers. They gave their written informed consent and received 30 euros at the end of the experiment. Ethical approval was obtained from the University of Trento Ethics Committee and conducted according to the Declaration of Helsinki.

### Method details

#### Stimuli and experimental procedure

The videos (visual speech) were recorded with a digital camera (VP-D15i; Samsung Electronics) at a rate of 25 frames per second. The audio files were recorded at a sampling rate of 44.1 kHz. The speakers were native Italians (two males and one female). Nine short text pieces (see supplementary material for examples) were recorded from each speaker lasting from 21 to 40 seconds each, resulting in 27 forward videos and audio files and, by reversing them, in 27 backward video and audio files. On average 36 pieces were randomly selected and presented to each participant counterbalancing the gender of the speakers. The mute videos were displayed on a projector panel in the MEG chamber and the audio files were presented binaurally via a sound pressure transducer (Etymotic Research ER-2) through two plastic tubes terminating in plastic earplugs while participants were fixating on a cross at the centre of the projector panel. The order of the visual and the auditory sessions was counterbalanced. Participants were instructed to passively watch the mute videos and listen to the audible speech. The experiment lasted less than one hour including preparation. Presentation was controlled via Psychtoolbox [32].

#### MEG recording

MEG was recorded at a sampling rate of 1 kHz using a 306-channel (204 first order planar gradiometers) VectorView MEG system (Elekta-Neuromag Ltd., Helsinki, Finland) in a magnetically shielded room (AK3B, Vakuumschmelze, Hanau, Germany). MEG signal was online high-pass and low-pass filtered at 0.1 Hz and 330 Hz respectively. Prior to the experiment, individual head shapes were digitized for each participant including fiducials (nasion, pre-auricular points) and around 300 points on the scalp using a Polhemus Fastrak digitizer (Polhemus, Vermont, USA). The head position relative to the MEG sensors was continuously controlled within a block through five head position indicator (HPI) coils (at frequencies: 293, 307, 314, 321 and 328 Hz). Head movements did not exceed 1.5 cm within and between blocks.

### Quantification and statistical analysis

#### Extraction of acoustic speech envelope and lip contour signal

The acoustic speech envelope was extracted using the Chimera toolbox by Delguette and colleagues (http://research.meei.harvard.edu/chimera/More.html) following a well-established approach in the field [8,33] where nine frequency bands in the range of 100 – 10000 Hz were constructed as equidistant on the cochlear map [34]. Sound stimuli were band-pass filtered (forward and reverse to avoid edge artifacts) in these bands using a 4^th^-order Butterworth filter. For each band, envelopes were calculated as absolute values of the Hilbert transform and were averaged across bands to obtain the full-band envelope that was used for coherence analysis. The envelope was then down-sampled to 256 Hz to match the down-sampled MEG signal (Fig. 1).

The lip contour of the visual speech was extracted with an in-house algorithm in MATLAB calculating the area (function *polyarea.m*) defined by eight points on the lips of the speaker (Fig. 1A). These points were defined in the first frame of the video and were tracked using the Kanade-Lucas-Tomasi algorithm (KLT, function *vision.PointTracker)* on the next frames [35,36]. The fluctuations of the mouth area are thus expressed in this signal at the sampling rate of the video (25 frames/sec), which was interpolated to 256 Hz to match the MEG signal.

#### MEG preprocessing

Data were analyzed offline using the Fieldtrip toolbox [37]. First, a high-pass filter at 1 Hz (6th order Butterworth IIR) was applied to continuous MEG data. Then, trials were defined keeping 2 sec prior to the beginning of each sentence and post-stimulus varying according to the duration of each sentence. Trials containing physiological or acquisition artifacts were rejected. Bad channels were excluded from the whole data set. Sensor space trials were projected into source space using linearly constrained minimum variance beamformer filters (van Veen et al., 1997) and further analysis was performed on the obtained time-series of each brain voxel. The procedure is described in detail here: http://www.fieldtriptoolbox.org/tutorial/shared/virtual_sensors. We used the standard template brain MRI from Montreal Neurological Institute (MNI) aligned in individual headspace. Personalized MRIs were created by warping the standard MRI to optimally match the individual fiducials and headshape landmarks [39]. A 3D grid covering the entire brain volume (resolution of 1 cm) was created based on the standard MNI template MRI. The MNI space equidistantly placed grid was then morphed to individual headspace. Finally, we used a mask to keep only the voxels corresponding to the grey matter (1457 voxels). Using a grid derived from the MNI template allowed us to average and compute statistics as each grid point in the warped grid belongs to the same brain region across participants, despite different head coordinates. The aligned brain volumes were further used to create single-sphere head models and lead field matrices [40]. The average covariance matrix, the head model and the leadfield matrix were used to calculate beamformer filters. The filters were subsequently multiplied with the sensor space trials resulting in single trial time-series in source space. For both power and coherence, only the post-stimulus period was considered and the initial long sentence-based trials were cut in trials of 2 seconds to increase the signal-to-noise ratio. The number of forward and backward trials was equalized. The number of forward and backward trials did not differ statistically for the female and male speakers.

#### Coherence calculation

The cross-spectral density between the acoustic speech envelope and the corresponding lip contour was calculated on single trials with multitaper frequency transformation (dpss taper: 1–20 Hz in 1 Hz steps, 1 Hz smoothing). The same was then done between each virtual sensor and the acoustic speech envelope as well as the lip contour (dpss taper; 1 – 130 Hz in 1 Hz steps; 3 Hz smoothing). Then, the coherence between activity at each virtual sensor and the acoustic speech envelope or the lip contour while participants either watched the lip movements or heard the speech was obtained in the frequency spectrum and averaged across trials. We will refer to the coherence between acoustic speech envelope and brain activity as acoustic speech-brain coherence and between the lip-contour and brain activity as lip-brain coherence.

#### Granger causality

We took all voxels of the statistical effect (Fig. 2A) between forward and backward unheard acoustic speech-brain coherence and from those identified for each participant an individual voxel based on the maximum difference in forward vs. backward unheard acoustic speech-brain coherence that occurred. This voxel was then used as a seed for calculating Granger causality during video presentation (visual speech) with all other voxels [41]. Fourier coefficients were calculated using multitaper frequency transformation with a spectral smoothing of 3 Hz followed by calculating bivariate granger causality. This led to measures of ingoing and outgoing connections for the individual seed voxel for forward and backward visual speech which we normalized ((ingoing − outgoing)/(ingoing+ outgoing)) separately for forward and backward visual speech (see also [42,43]). Note that in the following, whenever we speak of Granger causality we refer to normalized Granger causality with an individual occipital voxel as seed region.

#### Statistical analysis

To test for differences in source space, forward vs. backward contrast was performed for acoustic speech-brain coherence on a spectrum level discarding the time dimension for the whole brain. A two-tailed dependent samples t-test was carried out averaging the coherence values over our frequency band of interest, theta (4 – 7 Hz), as the lip-speech coherence showed a clear peak in this frequency. Consistent with Ghitza [6] and Giraud and Poeppel [7] in our stimulus material, this frequency corresponded with the frequency of syllables (mean=5.05 Hz, range: 4.1–5.6 Hz). Note that when contrasting the forward and backward coherence between MEG activity and the acoustic speech envelope watching lip movements no acoustic signal was present. For the Granger causality, we also averaged values over our frequency band of interest, theta (4 – 7 Hz). Given findings of enhanced top-down coupling during intelligible speech compared to unintelligible speech in the left hemisphere [25], we expected more connectivity during the forward condition (intelligible) and therefore used a one-tailed t-test as test statistic. Further, based on models of dual-route processing of speech [3,5] proposing mapping of acoustic signals within specific regions of the left hemisphere, we took a more statistically focused approach and accordingly selected regions in left temporal, parietal, postcentral regions and precentral regions as defined by AAL atlas implemented in fieldtrip including the regions of interest as proposed by the dual-stream model [5].

To control for multiple comparisons, a non-parametric Monte-Carlo randomization test was undertaken [44]. The t-test was repeated 5000 times on data shuffled across conditions and the largest t-value of a cluster coherent in time and frequency was kept in memory. The observed clusters were compared against the distribution obtained from the randomization procedure and were considered significant when their probability was below 5%. Significant clusters were identified in space. For contrasting forward and backward lip-speech coherence we used montecarlo permutation with FDR for multiple comparisons correction.

For statistical comparison of the individually extracted values of occipital forward vs. backward unheard acoustic speech-brain coherence and occipital lip-brain coherence, we used two-tailed t-tests. The results of the occipitally extracted forward vs. backward acoustic speech-brain coherence and (unseen) lip-brain coherence during actual acoustic speech can be found in the supplementary material.

We correlated the effects from the unheard acoustic speech-brain coherence with the results of the Granger causality analysis. We calculated the mean across voxels from the occipital coherence effect during the forward visual speech and correlated this with the selected regions in left temporal, parietal, postcentral regions and precentral regions. While a significant effect was found, we also report the effects above the statistical critical value that did not survive cluster-correction, as the patterns overlap strongly with the regions obtained from the Granger contrast, adding support to their potential functional relevance. We looked at the conjunction between the effects from the forward-backward Granger causality contrast and the correlation analysis. Voxels in precentral and postcentral gyrus showed overlapping effects, and for the voxel with the strongest correlation we display the scatterplot illustrating the correlations between significant voxel of the forward-backward unheard acoustic speech-brain coherence contrast and the overlapping voxels of the Granger causality forward-backward contrast and the correlation analysis.

For visualization, source localizations of significant results were mapped onto inflated cortices using Caret [45] or a standard MNI brain as implemented in Fieldtrip. Anatomical regions were labelled using the AFNI-atlas [46].

#### Behavioral experiment

To elucidate if visual speech presented without sound also differs in terms of intelligibility, we performed an independent behavioral experiment with 19 Italian native speakers (age 32.4 ± 3.9, 12 females). We used the same stimuli as in the MEG experiment cut in phrases of 3.5 – 7.5 sec duration. The video with the phrases (25 forward and 25 backward) were presented without sound and at the end of each trial, two words appeared on the screen and the participant responded by button press which of the two was contained in the short presented snippet. In both the forward and backward condition one option represented the correct word. Performance was statistically analyzed using t-tests between hit rates of the forward and backward condition and between chance level (50%) and the hit rates.

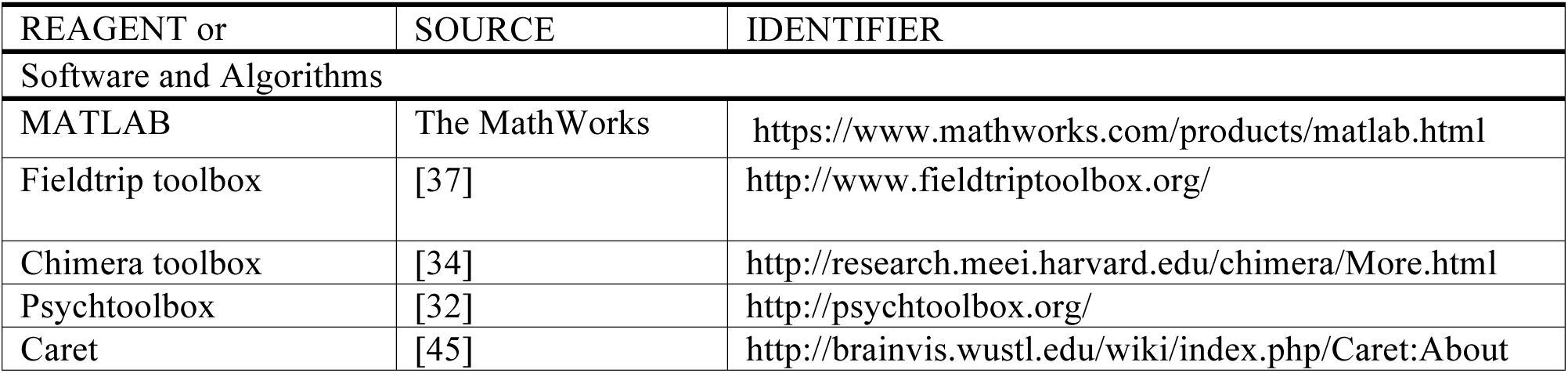
KEY RESOURCES TABLE

## Supplementary information

**Material:** text pieces used in MEG study

1. T01_s01: Mi telefona un tale per dirmi che sta facendo una piccola inchiesta. E vorrebbe che gli rispondessi a questa domanda: “Di che nazionalità vorrei essere se non fossi italiano”. Viviamo nel secolo delle domande. Chiudo gli occhi, aspiro profondamente e rispondo: “Prima di tutto, bisognerebbe provare che sono italiano. Non scrivo e non parlo il mio dialetto. Non adoro la città dove sono nato. Il gioco del calcio non mi entusiasma. Lo sopporterei se sul campo i giocatori fossero venti mila e il pubblico ventidue persone. Non ascolto la radio e non guardo la televisione.”
2. T01_s02: pago le contravvenzioni. Non ho amici negli uffici importanti e mi sarebbe penoso partecipare a un concorso. Non so cantare e non mi piace sentir cantare gli altri, se non a teatro. Non scrivo versi. Ho conservato sempre gli stessi amici. Mi piace viaggiare per l’Italia e quasi ogni luogo mi incanta e vorrei restarci. Sotto questo aspetto potrei essere un inglese.
3. T01_s03: Se visito un museo non parlo ad alta voce. E se vado in una biblioteca non tento di portarmi via un libro o le sue illustrazioni. Tuttavia, che io sia italiano potrebbe essere negabile. Infatti mi piace dormire, evitare le noie, lavorare poco, scherzare e ho un pessimo carattere, perlomeno nei miei riguardi.
4. T02_s01: “Prendiamo un caffè?” Significa “interrompiamo quello che stiamo facendo?”, “ti devo parlare.”, o anche “stiamo un po’ insieme”. Prendere un caffè è qualcosa che oltrepassa il caffè. Un esortazione a cambiare qualcosa, registro, dimensione, approccio. Se dite “sì, grazie, prendiamoci un caffè” accettate un invito più ampio. Cambiate luogo passando dal lavoro al bar. Cambiate la dimensione dei rapporti, dal formale all’ informale. Cambiate l’atmosfera.
5. T02_s02: Prendere un caffè modifica le distanze, muta i rapporti, avvicina. E’ un luogo denso di simboli, il caffè. E una volta arrivati al bar, il caffè è come se l’aveste già preso. Quindi potete tranquillamente bere qualcos’altro, un succo di frutta, un vino o un’acqua minerale con una fetta di limone. Se però decidete di andare fino in fondo e prendere davvero un café, vi si apre davanti un mare sconfinato di possibilità. E di nomi.
6. T03_s01: Per capire in quale conto il siciliano tiene la puntualità, basta considerare il modo in cui generalmente da un appuntamento. Egli non dirà per esempio “incontriamoci alle 9”, ma userà un’espressione ben più vaga e indefinita. “Ci vediamo verso le 9?” dove quel “verso” strategicamente collocato prima dell’ora stabilita autorizza implicitamente un ritardo di almeno mezzora. L’approssimazione infatti è sempre per eccesso mai per difetto. Chi dovesse arrivare un po’ prima delle 9, rientrerebbe incautamente in un verso che riguarda le 8.
7. T03_s02: Da noi non si gioca mai d’ anticipo. Posticipare, dilazionare, differire è la nostra arte. Dissipare il tempo, è una filosofia della vita, ancor che appena abbozzata che ha notevoli ricadute, quasi sempre negative, nelle attività economiche. L’artigiano o l’operaio che deve svolgere un lavoro a casa del cliente è solito dare un’indicazione orientativa del suo intervento. Verrà nel pomeriggio o in serata o in mattinata. Non gli potremo estorcere niente di più rassicurante.
8. T04_s01: Il viaggio è durato cinque anni. E la ragazza era all’ epoca ancora molto piccola. La cosa più importante per lei, oltre ad avere sempre con se il suo pupazzo orsetto, era che mamma e papa si prendessero cura di lei giorno e notte. A meno che non fossero impegnati in una riparazione o in una manovra difficile.
9. T05_s01: La qualità della vita, l’arte e la scienza di vivere bene, di spassarsela non hanno nessuna relazione con la qualità delle vacanze. Secondo me, vivere significa lavorare il giusto e bene, mantenersi in salute, avere facile accesso alle conoscenze, muovermi velocemente ed economicamente nella mia città e nel mio paese. Ed avere facilitati rapporti sociali. Poter scegliere tra molte opzioni come disporre del mio tempo libero in modo proficuo e creativo.

**Results:**

**Fig. 1S:**
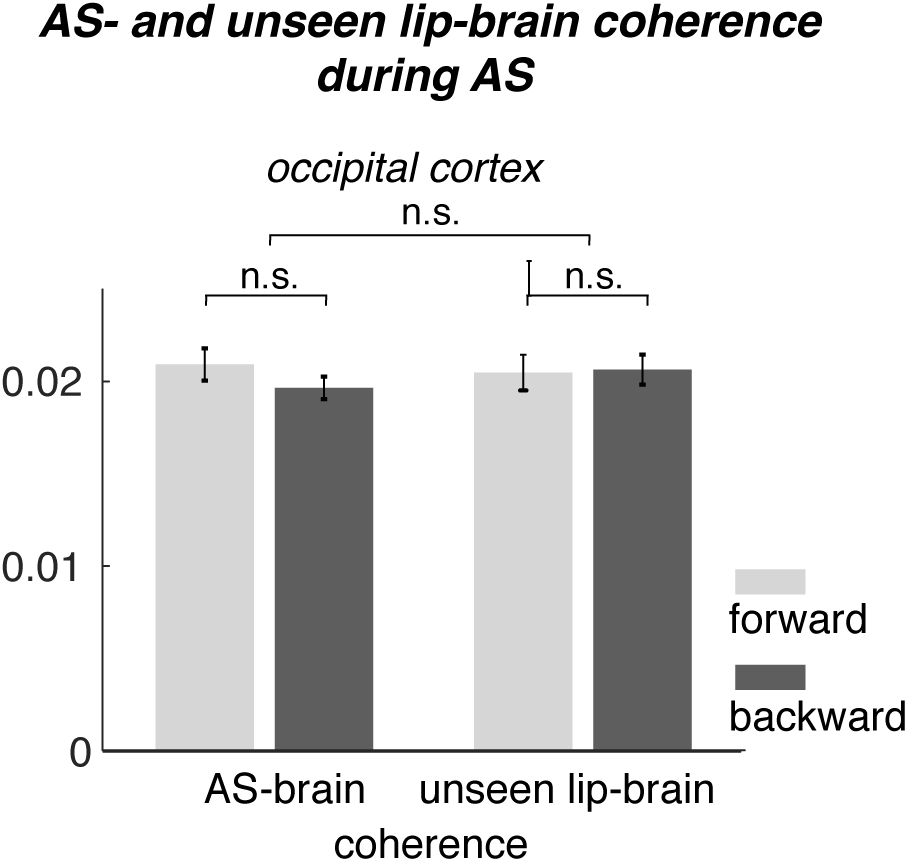
Mean of the individual acoustic speech (AS)-brain and unseen lip-brain coherence values (extracted at the occipital voxels of the statistical effect found for forward versus backward unheard acoustic speech-coherence) during acoustic speech. Contrast of forward-backward acoustic speech-brain coherence (t(23)=1.29, p=0.21) and unseen lip-brain coherence (t(23)=−0.12, p=0.9) did not show differences. Also, the contrast between forward-backward difference for acoustic speech-brain and unseen lip-brain was not significant (t(23)=0.88, p=0.39).

